# Non-canonical role for ATPase HSC70 in driving Clathrin remodeling during compensatory endocytosis in synapse

**DOI:** 10.1101/2025.04.02.646821

**Authors:** Sai Krishnan, Yaroslav Tsytsyura, Nataliya Glyvuk, Julia Lehrich, Jürgen Klingauf

## Abstract

Neurons use Clathrin-mediated endocytosis to retrieve synaptic vesicle (SV) proteins during compensatory endocytosis after presynaptic SV fusion. We have shown SV cargo to be re-sorted and pre-assembled outside the active zone into Clathrin-coated structures (CCS) of variable size and curvature, constituting a readily retrievable pool. During compensatory endocytosis CCS of the readily retrievable pool must be remodeled swiftly into spherical vesicles within 10 seconds at physiological temperature. How this is achieved remains elusive. Here we performed live-cell imaging on intact as well as scanning electron microscopy on unroofed hippocampal Xenapses, TIRFM-amenable presynaptic boutons formed en-face on microstructured and functionalized coverslips. While CCS can slowly remodel into spherical pits in unroofed Xenapses within tens of minutes, this process is highly accelerated in intact Xenapses, as evidenced by the rapid (< 10 seconds) exchange of EGFP-labelled Clathrin and AP2 adaptor after photo-bleaching. This fast remodeling of CCS was observed in the absence of stimulation and cannot be explained by constitutive endocytosis. Hence, this process must be driven by cytosolic factors which are lost during unroofing. Using membrane-permeant interfering peptides we identify Hsc70, the ATPase which along with Auxilin drives uncoating of endocytosed Clathrin-coated vesicles, to have an additional role in driving curvature of invaginating CCS.

## Introduction

The fundamental unit of Clathrin mediated endocytosis (CME) in its varied functions is the Clathrin-coated pit (CCP), Clathrin triskelia assembled into lattice structures at the plasma membrane, which then pinches off to generate Clathrin-coated vesicles ^1–4^. CME performs its endocytic function in conjunction with a range of accessory proteins and is a remarkably resilient and ubiquitous pathway ^5–7^. Furthermore, membrane-associated Clathrin cages being observed in unroofed cells ^8^ has led to the formulation of the readily retrievable pool (RRetP) ^9–13^, a pre-sorted and preassembled complex of Clathrin and associated proteins to enable the precise endocytosis of synaptic vesicles (SV).

This has made elucidating Clathrin mechanism at endocytic pits the focus of numerous studies wishing to identify its underlying properties. A key feature of Clathrin which may explain its functional significance is its dynamic properties, reflected in the ability of Clathrin to spontaneously curve into cages of varied sizes and shapes ^2,8^. The potential for the inherent plasticity of Clathrin as a contributor to changes in membrane curvature forms the basis of the long proposed constant surface area model of Clathrin organization ^8^. Indeed, live studies of CCPs using fluorescence recovery after photobleaching (FRAP) reveal high turnover rates with Clathrin recovery occurring within seconds of the photobleaching pulse suggesting significant remodeling of the surface Clathrin ^14–16^. The constant curvature model challenges the constant surface area model and the arguments against the latter mechanism rests on energetic unfavourability of the incorporation of pentagons into the predominantly hexagonal flat Clathrin lattice as it buckles and transitions to a curved pit ^17–20^. Other studies have sought to settle on a compromise model moving away from the extremes of the above two mechanisms. These posit a Clathrin pre-assembly of 50-70 % of its final surface area before the onset of curvature ^21–23^.

Thus, there exists in the Clathrin field, despite more than 5 decades of research, uncertainty regarding many aspects of its ability to organize and invaginate the nascent endocytic pit. A contributory factor to this is that most Clathrin studies have employed non-neuronal cells, which enable direct visualization of surface proteins by total internal fluorescence microscopy (TIRFM), but where endocytosis is constitutive and cannot be triggered. This vitiates the dynamic Clathrin exchange observed in fluorescence recovery after photobleaching (FRAP) assays, which provide the most promising *in vivo* evidence for Clathrin remodeling. The inaccessibility of the small presynapse has also made it challenging to conclusively demonstrate that surface assembled Clathrin, as part of the RRetP, is the primary unit of compensatory endocytosis.

In this study we addressed this bottleneck by investigating Clathrin remodeling and its underlying mechanism using our established TIRFM-amenable pre-synapse model, the Xenapse ^9^. We have previously demonstrated the unroofed Xenapse membrane surface to be studded with preassembled Clathrin demonstrated to constitute the RRetP with a defined endocytic capacity ^9^. Here, the morphologically diverse surface CCS in unroofed Xenapse displayed spontaneous ability to gradually form highly curved pits. This remarkable capacity for reorganization was reflected in the rapid recovery following photobleaching of the surface Clathrin under physiological conditions and with minimal endocytic contribution. Finally, by acute treatment with membrane permeable Hsc70-binding peptide we were able to significantly reduce the rate of surface Clathrin exchange and were able to demonstrate a non-canonical role for Hsc70 in maintaining the RRetP for compensatory endocytosis. Recent studies have argued against the salience of CME in the pre-synapse on the basis of it not being fast enough for cargo retrieval ^24–26^. The ability of pre-synaptic surface Clathrin to rapidly remodel under physiological conditions, in a process aided by additional actors such as Hsc70, argues against CME being too unwieldy to re-establish presynaptic homeostasis.

## Results

### Spontaneous curvature of Clathrin coated structures in unroofed Xenapse shows their intrinsic flexibility

The surface micropatterning of the coverslip required for Xenapse formation results in the boutons growing in an organized pattern on the coverslip surface (Fig. 1a). In addition to making them TIRFM-amenable, the patterning lends them to being unroofed by the application of a short sonicator pulse, a technique pioneered by Vacquier and Clarke ^27,28^ and later extensively employed by John Heuser to study surface Clathrin structures in non-neuronal cells ^8,29^. Briefly, the apical membrane of the intact Xenapses is removed i.e. unroofed in nominally 0 Ca^2+^ buffer followed by fixation and processing for scanning electron microscopy imaging (Fig.S1a and b). In our previous publication, by applying the unroofing approach to our model culture, we showed the surface Clathrin coated structures (CCS) to constitute the RRetP, which defines the endocytic capacity of the Xenapse ^9^. Here, we use the same technical approaches used in our previous studies to investigate a fundamental property of Clathrin, its capacity for remodeling, and how this impacts the endocytic function of the RRetP.

**Fig. 1:**
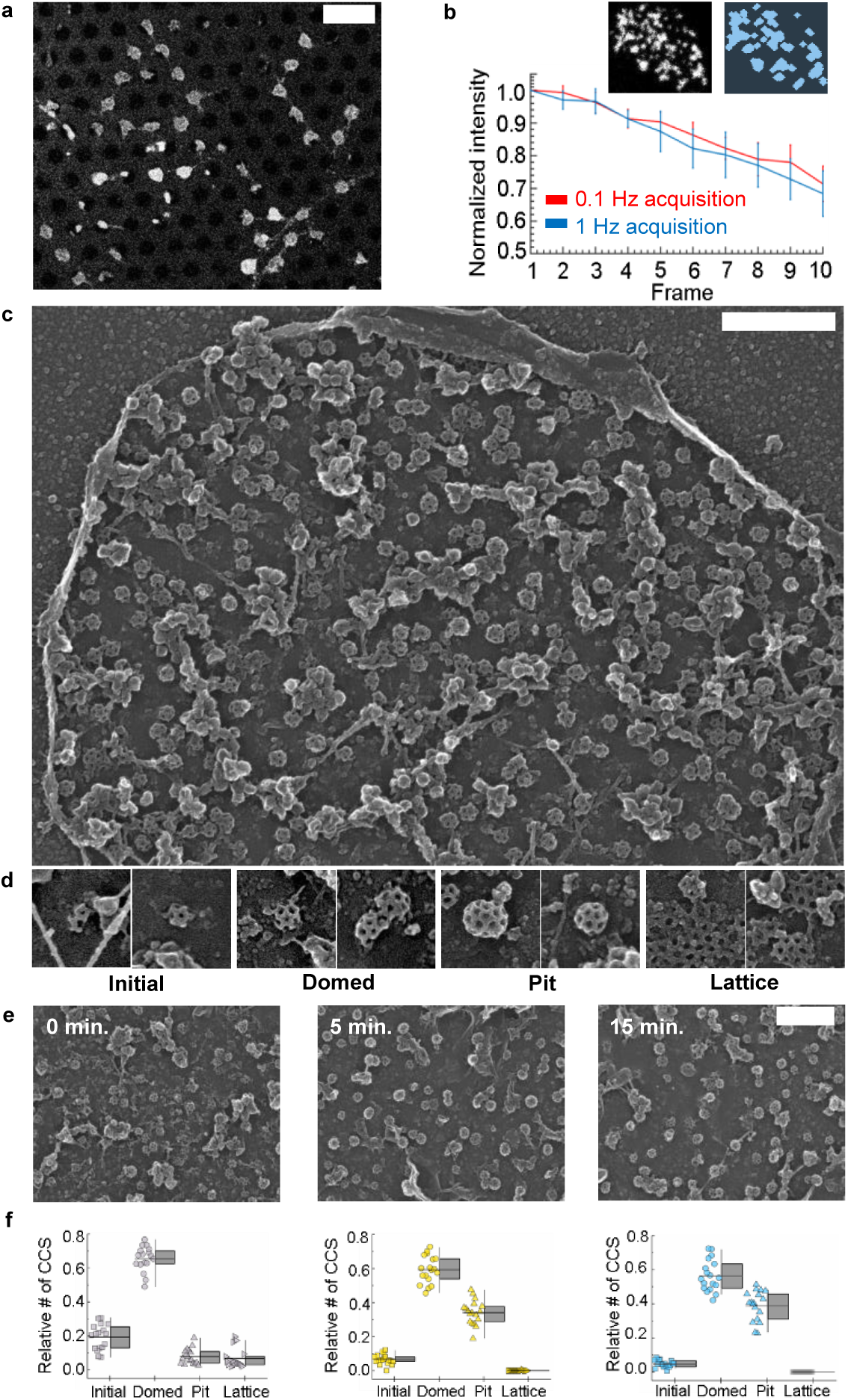
Preassembled Clathrin on the xenapse surface exhibits an intrinsic capacity for rearrangement. **a**, Confocal overview of Halo-CLCa expressing and JF646 stained Xenapses. Scale bar 15 µm. **b**, Top: Example of the average image of a STED image stack of an unroofed Xenapse expressing Halo-CLCa (stained with JF646) (left) and its thresholding derived binary image (right) defining single CCS. Below: Average normalized fluorescence traces of single CCS observed above imaged at RT in freshly unroofed Xenapses. Same STED imaging parameters were used for measurements at 1 Hz (blue) and 0.1 Hz (red) measurements. The same level of bleaching between the traces indicates no loss of surface clathrin during the prolonged incubation (mean ± s.e.m. n = 7 and n = 5 boutons for 0.1 Hz and 1 Hz respectively from a single biological replicate consisting of >40 localized Clathrin structures). **c**, SEM image (an overview) of unroofed and immediately fixed Xenapse. Scale bar 500 nm. **d**, A gallery of the crops from SEM images of unroofed and immediately fixed xenapses showing the different stages of Clathrin coat formation. Scale bar 100 nm. **e,f**, Representative individual fragments from SEM images of xenapses fixed at 0 (left), 5 (middle), and 15 (right) min. after unroofing, used for quantification of different assembly states of CCS (bottom panels), shown as a fraction of the total number of CCS at corresponding time points. The combined graphs show the scatter plots of the data used to build the box charts. Horizontal lines in the box charts represent the median with interquartile ranges (boxes) and vertical lines show a 1-99% spread (n = 17 Xenapses for 0 and n = 18 Xenapses for 5 min. and 15 min.). Scale bar 500 nm.

We first established, whether the CCS on the unroofed surface are not destabilized and lost over time. STED imaging at 0.1 Hz and 1 Hz of Xenapses expressing the Halo-tagged neuronal Clathrin light chain a1 (Halo-CLCa, labelled with JF646) revealed single structures which gradually decrease in intensity (Fig. 1b). Photobleaching and not loss of surface Clathrin explains the similar extent of decline between shorter and longer acquisition durations, as the Halo-CLCa is imaged at the same duty cycle. Thus, any change in the CCS assembly we observed in unroofed preparations following prolonged incubation must be due to rearrangement of assembled membrane-bound Clathrin. Fixing and processing the Xenapses at various time points after unroofing replicated the result observed in non-neuronal cells ^30^ with the slow but spontaneously curvature of the surface CCS into highly curved pits evidently without any input from additional cytosolic factors (Fig. 1e and f). The initial units of Clathrin form a small fraction and drop slightly at longer time points suggesting minor loss or some contribution to the formation of the two larger structures. It is striking that the flat lattice structures so common in non-neuronal cells ^31,32^ are hardly to be observed in Xenapses. Despite the significant change in curvature between domed and pit CCS, the coat area and diameter are not much different (Extended Fig.1c-f). This shows the assembly levels of the domed and pit structures are not significantly different and change little over time. These results are consistent with the claimed propensity of surface Clathrin to constantly experience spontaneous fluctuations, which loosen but do not destabilize the assembled Clathrin ^33^.

### The preassembled surface Clathrin is dynamic and exhibits continuous remodeling at steady state

A significant upshot of the CCS existing in a plethora of different sizes with an inherent ability to spontaneously curve (Fig. 1c) is, that the surface Clathrin is prone to significant remodeling under physiological conditions. The standard approach to test the inherent stability of a complex is to perform FRAP experiments on the labelled structures where a reasonably rapid recovery in fluorescence will suggest the underlying protein is undergoing cycles of assembly and disassembly.

The surface EGFP-CLCa was observed in patches and the diffusion component of the post-bleach recovery was subtracted to reveal the main sites of binding exchange (Fig. 2a inset). The fluorescent recovery at these patches after an EW-FRAP pulse lasting 2 seconds showed a steady rapid recovery with a distinctly faster recovery at 37°C (Fig. 2ab).

**Fig. 2:**
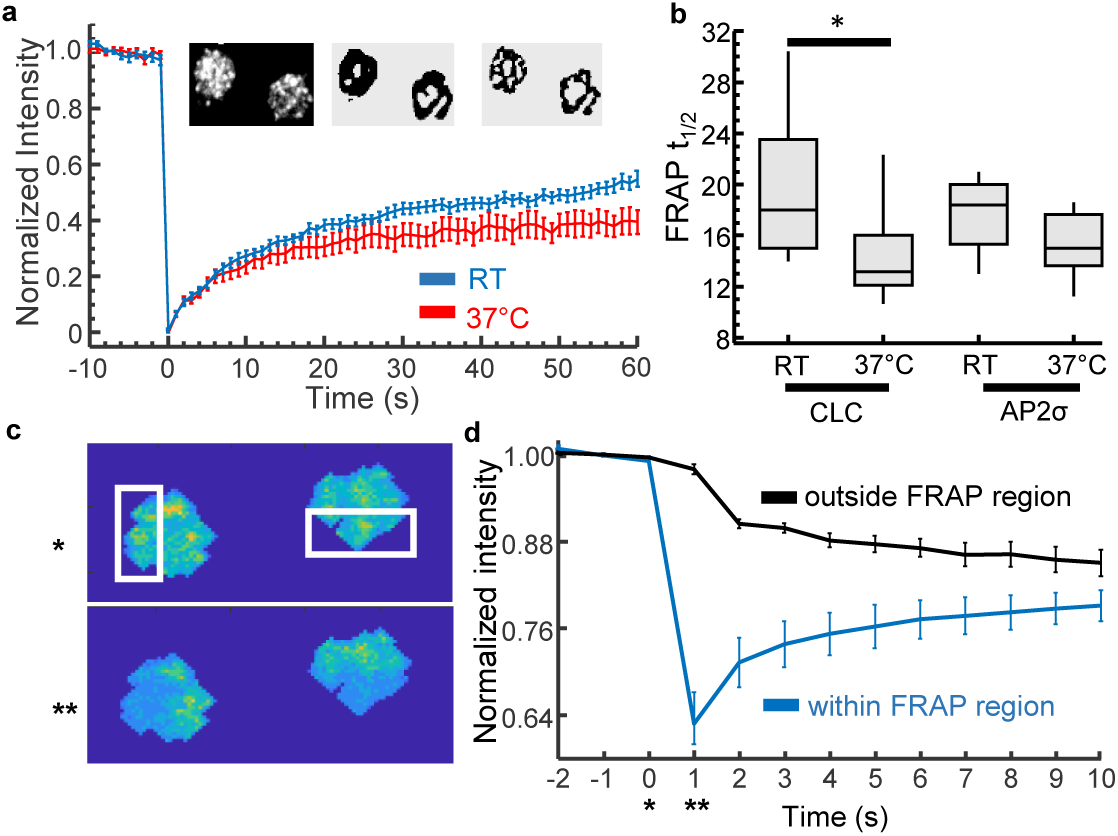
FRAP reveals dynamic exchange of preassembled surface Clathrin with the cytosolic pool. **a**, Normalized average of EGFP-CLCa fluorescence recovery traces at RT (n = 9 measurements) and 37°C (n = 13 measurements) in response to a TIRF photobleaching pulse revealing the amplitude of recovery (mean ± s.e.m. with data from 3 biological replicates). Inset: Representative TIRFM image of xenapses expressing EGFP-CLCa (left) with its à trous wavelet-filtered binary image (centre) and diffusion-component removed subregions (right), of which the latter was used to extract the FRAP response curves. **b**, Half-time of FRAP, i.e. times required for 50 % fluorescence recovery, for EGFP-CLCa at RT and 37°C and AP2σ-EGFP (Extended Figure 2a). Data shown as Boxplot with the median and interquartile ranges. *p < 0.05 by two sample Kolmogorov-Smirnov test. **c**, Pixel íntensity images of confocal fluorescence image of live xenapses expressing EGFP-CLCa with the frame before (above) and after (below) FRAP pulse taken at time 0 and 1 second in (**d)** respectively. The marked area indicates the xenapse region subjected to photobleaching pulse. **d**, Average normalized fluorescence intensity traces of pixels within and outside the Xenapse FRAP regions (n = 10 measurements consisting of 40 Xenapses in total from 3 biological replicates).

In contrast no such temperature dependent effect was observed for EGFP-AP2σ (Extended Fig.2a, Fig.2b), one of the subunits of the heteromeric adaptor protein AP2 which associates Clathrin to the plasma membrane lipid surface ^34^. The lack of temperature difference is likely due to the higher affinity of the adaptor complex to the lipid membrane. However, the very fact of not just Clathrin but also its adaptor displaying rapid recovery denotes the RRetP as a whole to be remodeling steadily and substantially in the absence of a stimulus. In light of the varied curvature of the surface CCS and their demonstrable tendency to reorganize (Fig.1), the FRAP exchange likely reflects the changes in curvature of the surface CCS and makes the arguments about energetic constraints involved in Clathrin remodeling redundant.

As a control, the scission protein DynaminI-EGFP was tested for recovery and the FRAP response did not reveal a membrane binding component but only a diffusional one (Extended Fig.2b). This is consistent with Dynamin being recruited after stimulation ^35^, and not being preassembled as has been claimed in previous publications ^25^. It is also notable that the post-diffusion recovery converges with the binding component recovery of EGFP-CLCa. This is significant as it suggests the lack of full Clathrin recovery to the baseline is due to a portion of cytosolic EGFP-CLCa being bleached during the 2-second FRAP pulse and not due to the presence of an immobile fraction, yet again implying a substantial remodeling of the Clathrin RRetP. Does this exchange only occur at the edges of CCS or across the whole coat area? While technical limitation prevents answering this question at the single CCS level, we addressed this by bleaching half of the Xenapse in confocal mode (Fig. 2c). The resulting fluorescence recovery in the bleached half very nearly converges with the loss of fluorescence in the unbleached sector of the same Xenapses (Fig. 2d). This convergence implies, that exchange of triskelia and AP2 indeed occurs across the whole CCS area. Finally, we used photoswitchable mEOS3.2-CLCa to convert the surface Clathrin to red fluorescence. This revealed not just a steady recruitment of the non-converted cytosolic mEOS3.2-CLCa to CCS to the membrane (Extended Fig.2c) but also removal of converted mEOS3.2-CLCa from the surface (Extended Fig.2d).

### The dynamic Clathrin exchange at steady state underlies the substantial stimulus-dependent remodeling of surface CCS

Overexpressed Clathrin labels the surface pits in the evanescent field and electrical stimulation leads to endocytic removal followed by slow recovery, reflecting the reformation of the RRetP out of newly exocytosed SV proteins, adaptor proteins and Clathrin for subsequent rounds of endocytosis ^9^ (Fig. 3a). A simple difference image of the endocytic minimum (Fig. 3a II) with the baseline (20 seconds before stimulus, Fig. 3a I) and back-baseline (final 10 seconds, Fig. 3a III) can highlight the regions of endocytosis and reformation respectively. The surface Clathrin is densely present with the TIRFM image of EGFP-CLCa showing non-uniform localization of CCS (Fig. 3b I). On stimulus, a fraction of the surface structures is retrieved and reformed but in only partially overlapping sub-regions (Fig. 3b II, III and composite). The minimal overlap between the two sub-regions suggests a substantial degree of stimulus dependent reorganization of the surface CCS. Focus on the pre-stimulus basal pit dynamics at the single CCS level using STED nanoscopy reveals only a small subset of CCS undergoing spontaneous endocytosis or being reformed in a 20 second interval (Fig. 3c and d). These STED nanoscopy images show most of the surface CCS are positionally quite stable over tens of seconds (Fig. 3e). Furthermore, the overexpressed Halo-CLCa structures almost precisely match the endogenous Clathrin structures stained by antibodies in unroofed Xenapses demonstrating the Halo-CLCa to faithfully incorporate into endogenous structures (Fig. 3f).

**Fig 3:**
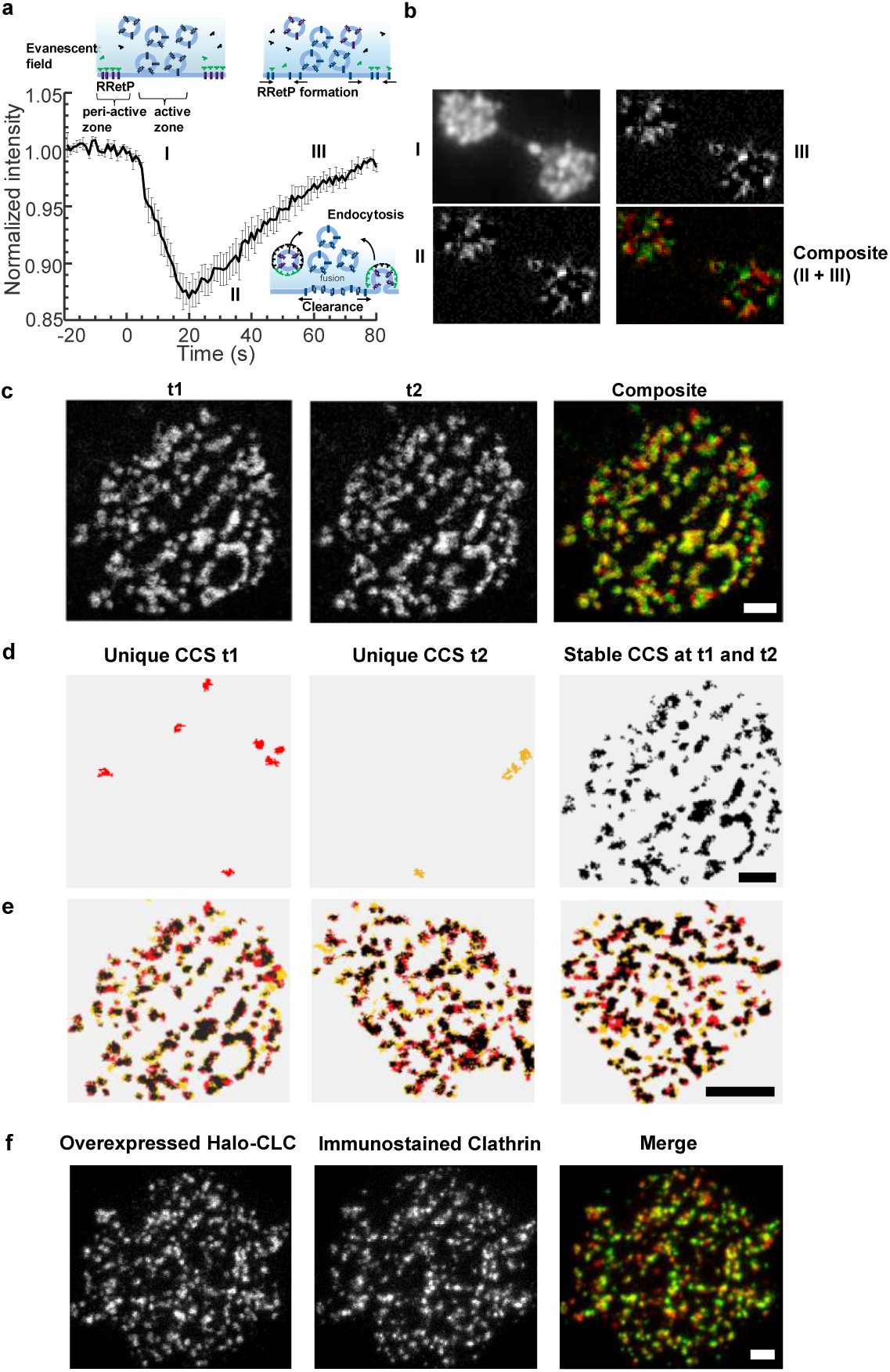
Stimulation elicits substantial remodeling of surface Clathrin which remains stable at steady state. **a**, Average fluorescence trace at RT of surface EGFP-CLCa in Xenapses in response to 50 AP at 40 Hz (n = 8 measurements from 3 biological replicates). Inset schematic shows the EGFP-labelled surface Clathrin within the evanescent field before, during and after endocytosis. **b**, TIRFM image of Xenapse showing surface EGFP-CLCa, before stimulus pulse, with its localization within sub-regions (I). II and III are difference images of these xenapses revealing the sub-regions of endocytosis and post-endocytosis recovery respectively. The composite image shows the endocytic decline (red) and post-endocytic recovery (green) to only partially overlap. Scale bar 1 µm. **c**, Time lapse STED images (t1 and t2, with the latter imaged 20 seconds later) of live Xenapse expressing Halo-CLCa and stained with JF 646. Scale bar represents 500 nm. **d**, Binary image generated by thresholding showing the Clathrin sub-structures unique to Xenapse at t1 (red), 20 seconds later (t2, yellow) and the stable CCS common at both time points (black). Below are the dynamic CCS unique to t1 and t2 and the composite of all the Clathrin structures (representative example of 5 Xenapses from two biological replicates). Scale bar represents 500 nm. **e**, Thresholding derived binary image of time lapse STED images (20 seconds apart) of 3 live Xenapses expressing Halo-CLCa (JF646) showing the composite of all the Clathrin structures. The first Xenapse is the composite of the panels in **d**. The stable CCS common at both time points are in black along with sub-structures unique to the first image (red) and the second image 20 seconds later (yellow). Scale bar represents 1 µm. **f**, STED nanoscopy of surface Clathrin in unroofed xenapse by simultaneous visualization of overexpressed Halo-CLCa (stained with JF646) and immunostained endogenous Clathrin heavy chain (n=6 xenapses from two biological replicates). Scale bar equals 500 nm.

The significance of these observations is two-fold. First, this confirms that in Xenapses unlike in non-neuronal cells only a minute fraction of the membrane CCS is undergoing spontaneous endocytosis in a 20 s interval and thus do not significantly contribute to the FRAP responses in Fig. 2a which thus purely reflect the unbinding rates of surface Clathrin. Secondly, the positional stability of the CCS at rest hides the steady exchange dynamics apparent in the FRAP response but contributes to the CCS remaining supple and flexible in order to reorganize in response to a stimulus.

### The ATPase Hsc70 enables Clathrin remodeling at the Xenapse surface

The FRAP data reveal the accelerated ability of surface Clathrin to remodel itself and strongly points to the presence of a Clathrin binding protein catalyzing this process *in vivo*. The ATPase Hsc70 has been postulated to be the candidate protein by chaperoning Clathrin and promoting its exchange ^36–39^, a membranal counterpart to its established function in depolymerizing post-endocytic Clathrin-coated vesicles ^40–42^. Here we sought to deploy a novel approach of acutely targeting endogenous Hsc70 in the Xenapse preparation using the cell-permeant TAT peptide conjugated to an Hsc70 binding sequence. In addition to the established FYQLALT synthetic sequence (TAT-optimal) ^43^ we also used the endogenous C-terminal tail sequence of the Clathrin heavy chain (TAT-QLMLT), demonstrated to be the unique Hsc70 binding site (Fig. 4a and b) ^42^. The efficacy of the former sequence can be tested by introducing negatively charged amino acids shown to reduce binding affinity to Clathrin (TAT-optimal mutant) ^44^. TAT-QLMLT being longer than TAT-optimal renders it amenable to be scrambled and the resulting control peptide (TAT-scrambled) has the advantage of maintaining the same membrane penetrant properties.

**Fig. 4:**
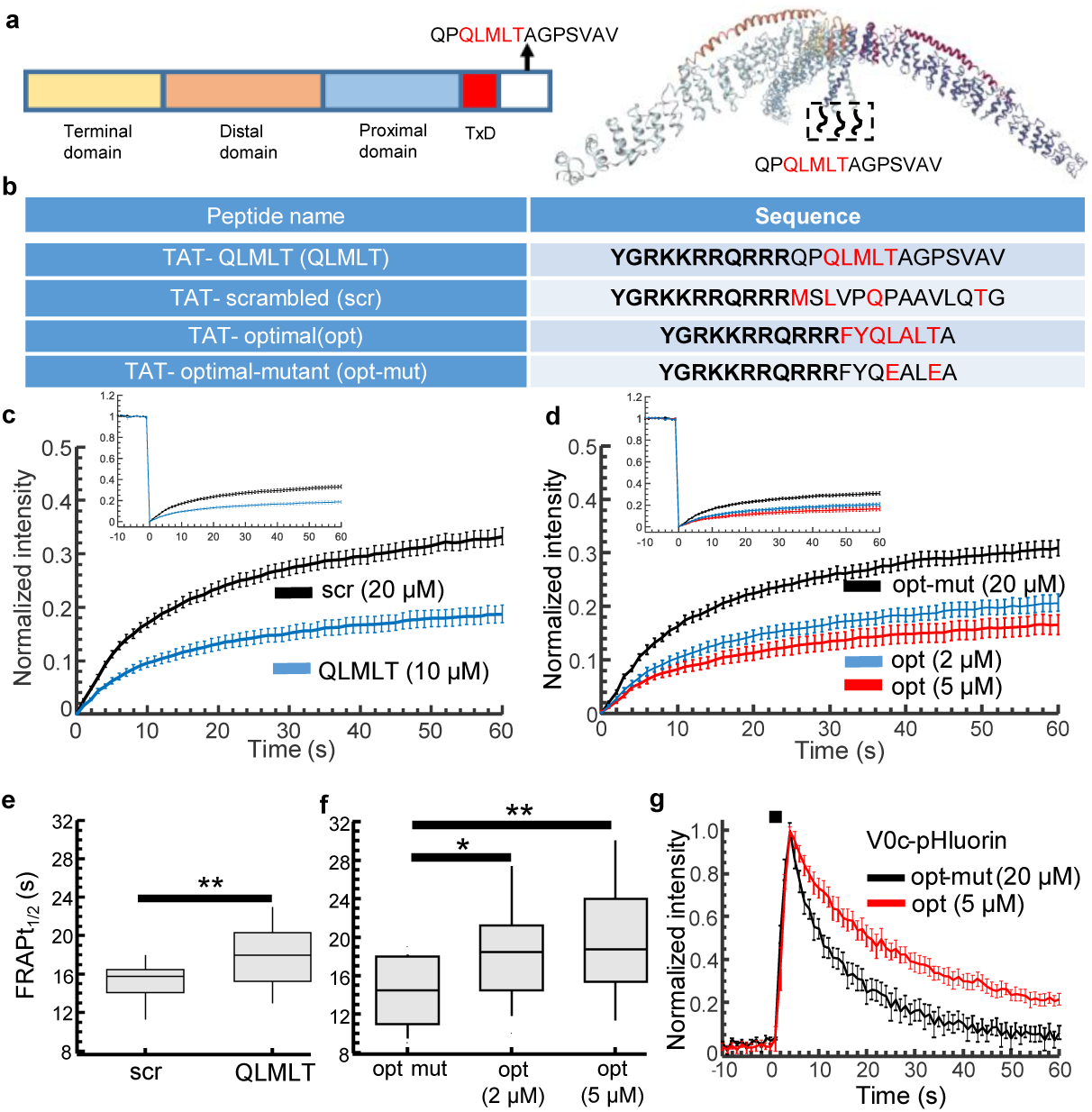
Rapid exchange of surface Clathrin catalyzed by Hsc70 is important for endocytosis. **a**, Clathrin heavy chain domain structure (left) and the crystal structure of Clathrin triskelia, the heavy chain and light chain complex (right, PDB ID: 3LVH) both showing the location of the Hsc70 binding hydrophobic sequence, QLMLT. **b**, Amino acid sequence of TAT peptides with common TAT sequence (black and bold) and the previously identified Hsc70 binding residues (red). TAT-QLMLT is the endogenous clathrin heavy chain C-terminal sequence containing the characterized Hsc70 binding motif. The endogenous sequence was rearranged to create the TAT-scrambled control peptide. TAT-optimal is the synthetic sequence optimized for high affinity binding to Hsc70. This affinity is significantly reduced by the replacing key residues with the negatively charged glutamate (TAT-optimal mutant). **c**, Normalized average of EGFP-CLCa fluorescence recovery traces at 37°C in response to treatment with TAT-scrambled (scr 20 µM, n = 25 measurements) and TAT-QLMLT (QLMLT 10 µM, n = 27 measurements) in response to a TIRF photobleaching pulse revealing the amplitude of recovery. **d**, Normalized average of EGFP-CLCa fluorescence recovery traces at 37°C in response to treatment with TAT-optimal mutant (opt mut 20 µM, n = 15 measurements) and TAT-optimal (opt 5 µM and 2 µM, both n = 18 measurements) in response to a TIRF photobleaching pulse revealing the amplitude of recovery. Insets in (**c**) and (**d**) showing the same recovery traces with the pre-bleach baseline included (mean ± s.e.m. from 3-5 biological replicates). **e,f**, Half-times of FRAP for data in (**c**) and (**d**) shown in (**e**) and (**f**) respectively. Data shown as Boxplot with the median and interquartile ranges. *p < 0.05 and **p < 0.01 by two sample Kolmogorov-Smirnov test. **g**, Normalized average of Voc-pHluorin transduced Xenapses responding to 100 AP at 40 Hz in the presence of TAT-optimal mutant (opt-mut 20 µM) and TAT-optimal (opt 5 µM) (mean ± s.e.m. n=7 measurements from 3 biological replicates).

The Clathrin FRAP response in the presence of the active peptides shows a significant decrease in the extent of Clathrin undergoing dynamic exchange (Fig. 4c and d) as well as the rate at which it does (Fig. 4e and f) relative to the respective control peptides. The change in the amplitude of recovery confirms the role of Hsc70 and its cofactors in chaperoning the freely floating Clathrin triskelia ^45–47^. The peptides, by sequestering the endogenous Hsc70, impair the dynamic exchange of the Clathrin units reflected in rate of surface Clathrin exchange being dampened. As the Clathrin dynamics are closely linked to changes in the assembly of surface CCS, it stands to reason that Hsc70 could catalyze changes in Clathrin *in vivo*, which we show to occur at a slower rate in the absence of Hsc70 (Fig. 1e). Finally, we tested the TAT-opt, the more potent of the two Hsc70 binders, on Voc-pHluorin expressing Xenapses. Upon stimulation with 100 APs endocytosis was significantly slowed and incomplete for TAT-opt compared to the low affinity TAT-opt mut (Fig. 4e). Notably, the control peptide (TAT-opt mut) as well as TAT-opt condition displayed normal exocytosis indicating these peptides do not disrupt membrane properties at the concentrations used. In summary, the inhibition of Hsc70-mediated Clathrin exchange has a significant effect on the SV recycling properties. Based on these findings we have come up with a new model of RRetP organization at the presynapse membrane surface summarized as a schematic (Extended Fig.3).

## Discussion

The organization of Clathrin on the membrane surface has been the topic of intense research but has been restricted to non-neuronal cells (PMID: 6987244, PMID: 2563729, PMID: 33823128, PMID: 28346440). The development of TIRFM-amenable presynaptic culture, the Xenapse, has allowed us here to directly investigate the properties underpinning the pre-synaptic Clathrin, which we have demonstrated to constitute the preassembled RRetP, the fundamental unit of compensatory endocytosis ^9^. The pre-synaptic CCP were revealed to be highly dynamic and susceptible to substantial changes in their assembly state in the absence of stimulation. We went on to reveal the mechanism underlying this remarkable plasticity with a non-canonical role for the ATPase Hsc70 in mediating the constant exchange of surface Clathrin.

The major cause for uncertainty over the nature of surface Clathrin organization lies in the supposed energetic unfavourability of changes in assembled Clathrin curvature as proposed by the constant surface area model. The deformation of hexagonal flat Clathrin structures into curved pits cannot occur without the introduction of lattice defects with the incorporation of pentagons ^48,49^. The constant curvature model purports to address this problem by proposing simultaneous changes in Clathrin assembly and membrane curvature ^18^. Evidence in favor of the constant area model of Clathrin organization has thus far largely relied on unphysiological conditions. Initial studies showed the formation of empty Clathrin microcages at the edges of Clathrin network under acidic conditions ^29^. In minimally reconstituted cell-free system Clathrin assembly alone appears to be sufficient to drive membrane invagination and spherical bud formation ^50^. Our data by demonstrating the capacity of shallow domed structure to curve into pits and by combining this with physiological data showing substantial Clathrin exchange in the absence of stimulation and endocytic activity, supports the growing evidence in favor of a cooperative model of surface Clathrin assembly ^23,51^. The consensus points to FBAR and adaptor proteins promoting the initial curvature and assembly, which initiates a feedback loop for Clathrin recruitment. This supports the notion of Clathrin having a role in sensing curvature ^52^. The missing part of the puzzle is whether a Clathrin chaperone exists to promote this transition at endocytic sites.

Attempts to demonstrate the role of ATP hydrolysis in Clathrin polymerization driven pit invagination has implicated the ATPase Hsc70, in addition to its established role in uncoating the Clathrin coated vesicle following fission, acting as a catalyzer of Clathrin mediated curvature ^38,39^. These studies however rely on non-neuronal cells where endocytosis is not triggered thereby making it difficult to distinguish the dynamic Clathrin exchange occurring at steady state. The strength of the findings in this study lies in being performed under physiological conditions and in neuronal preparations which allow for direct visualization of Clathrin without the aforementioned problems of non-neuronal cell lines. This makes the FRAP data showing the rapid exchange of not just surface Clathrin but also its adaptor AP2 more reliable and points to a substantial reorganization of the RRetP. Our data demonstrating the role of Hsc70 as a surface Clathrin chaperone makes the need for constant curvature redundant especially as Hsc70 catalyzes what we confirm to be the inherent ability of domed Clathrin to curve into pits by exploiting the potential energy loaded in the flat lattice. This remarkable malleability has been given structural credence by cryo-EM studies which reveal conserved but flexible contact points between Clathrin triskelia allowing fewer constraints in the deformation capacity of the assembled lattice structure ^53^. The relatively flat CCPs are thus not endocytic dead ends but unroofing captures them at early stages of their assembly/curvature cycle, which is attenuated by interfering with the Hsc70-Clathrin interaction. By perturbing this interaction acutely, using peptides characterized for their specificity and affinity ^42,43^, we avoided the pitfalls of previous attempts at addressing the role of Hsc70 at the membrane surface ^36,37^.

A shortcoming of this study is that we were unable to look at the Hsc70-mediated effect at a single CCS level or observe any structural changes to the surface Clathrin distribution in the presence of the Hsc70-sequestering peptide (data not shown). The latter effect was not observable as the slowdown in Clathrin exchange is not drastic at the concentrations of peptides used. At higher concentrations (above 5 or 10 µM), however, at which structural results would be observable for single CCS, the penetrant TAT peptide would cause membrane instability (measured with patch-clamp, data not shown). Another obstacle to observing Hsc70 at single pit level would be the low stoichiometry of interaction. *In vivo* studies have shown approximately 1 Hsc70 molecule per 6 Clathrin trimers for the uncoating of post-endocytic Clathrin coated vesicles when a maximal rate of disassembly is needed ^54^. At the lower disassembly rate at the membrane the number of Hsc70 molecules per Clathrin triskelia can be expected to be even smaller and challenging to detect in live assays.

This novel role for Hsc70 is likely to be universal across cell types but its specialized role in the presynapse is supported by the presence of the neuron-specific co-focator Auxilin I, which has been established to more potently catalyze Hsc70-mediated Clathrin disassembly in comparison to its ubiquitous homologue cyclin-G-dependent kinase (GAK) ^46^. The existence of such a dedicated mechanism in keeping the CCP supple and malleable is consistent with the need for the presynapse to respond rapidly and precisely in response to vesicle fusion. The Hsc70-mediated Clathrin exchange greatly aids the RRetP to rapidly replenish the pool of fusion competent SVs. It is notable that perturbing the Hsc70 interaction is detrimental to endocytosis and likely results from slower Clathrin exchange causing improper pit invagination (Fig. 4g). Hsc70-Clathrin may also have a role in sculpting the growing CCS, as suggested by the uniformly sized Clathrin pits in Xenapse (Fig. 1c), potentially linked to the need for precise packaging of SV cargo. The dynamic exchange of AP2 (Extended Data Fig. 2a), with its established cargo sorting role ^55–57^, lends credence to the idea that Hsc70 action on the CCP acts as a proofreading step for cargo packaging into the vesicle ^39^ and interfering with this would also be to the detriment of compensatory endocytosis.

## Materials and Methods

### Fusion constructs

The construction of EGFP-CLCa, expression construct with neuron specific Clathrin light chain (LCa1), has been described in previous publication ^12,15^. It was used to generate the Halo-CLCa by replacing the EGFP as also described in a previous publication ^9^. mEos3.2-C1 (Addgene plasmid # 54550) and psigma2-EGFP (Addgene plasmid # 53610) were obtained from Addgene (from the labs of Michael Davidson & Tao Xu and Tom Kirchhausen respectively). The CLCa from EGFP-CLCa was subcloned into mEos3.2-C1 to generate mEos3.2-CLCa. Dynamin1-GFP was courtesy of Prof. V. Haucke. Maxi-prep (Qiagen) purified constructs were used to transfect cells 4 days in vitro using a modified version of previously described Ca^2+^ Phosphate transfection protocol ^11^.

### TAT Peptide design and application

Four peptides encoding the TAT sequence (YGRKKRRQRRR) followed by either the endogenously derived (TAT-QLMLT) or synthetic (TAT-optimal) Hsc70 binding amino acid sequences and their respective non-binding variants (TAT-scrambled and TAT optimal mutant) were synthesized by GenScript Biotech (Netherlands) B.V. with >95% purity. Peptides were dissolved in deionized water and their stock concentrations determined using their absorbance and extinction coefficient at 280 nm. Peptide solutions were stored at −20°C. Before measurement, peptides were diluted to desired concentration in measuring buffer (140 mM NaCl, 2.4 mM KCL, 2.5 mM CaCl_2_, 1.3 mM MgCl_2_, 10 mM HEPES, and 10 mM D-Glucose in deionized water, pH adjusted to 7.4.) and applied onto the cells for 15 minutes incubation at 37°C. The measurement was performed in the presence of the peptide for no longer than 20 minutes.

### Animals

Breeding of C57BL/6N mice was conducted at the Central Animal Experimental Facility of the University Hospital Muenster according to the German Animal Welfare guidelines (permit number AZ81–02.05.50.20.019, issued by ‘Landesamt für Natur, Umwelt und Verbraucherschutz NRW’, Duesseldorf, Germany). Both male and female mice were used for primary neuronal cultures.

### Cell culture for Xenapse preparation

Primary cultures of hippocampal neurons from newborn (P0) C57/BL6 mice were seeded onto functionalized 18 mm micropatterned glass coverslips functionalized with Neuroligin 1(+AΔB) and placed in a Banker-type culture as described previously ^9^. Briefly, PDMS stamps with customized micropatterned surface corresponding to the Xenapse dimensions, were inked with PLL-PEG-HTL (PLL grafted with PEG-HTL). Micropatterning was achieved by bringing the inked stamps into contact with the plasma cleaned glass coverslip surface. The Neurexin endogenous to the hippocampal neurons are able to bind the surface immobilized Neuroligin 1 and thus result in the formation of presynaptic Xenapse culture.

Cells were grown on functionalized coverslips in Neurobasal-A medium supplemented with 2% B-27, 1% Glutamax, and 1% penicillin/streptomycin (all Thermo Fisher, MA, USA) under 5% CO_2_ at 37°C, for 16-17 days. All imaging was carried out at 14-18 days in vitro.

### Scanning electron microscopy of unroofed Xenapses

Unroofing of Xenapses at DIV 16-18 was performed according to previously published studies ^58,59^. In brief, cells were washed off the culture medium with calcium-free HEPES-buffered Ringer solution (in mM: NaCl 140, KCl 2.4, MgCl_2_ 4, HEPES 10, D-glucose 10, pH 7.3), then incubated for 15 seconds in the same solution containing 0.1 mg/ml PLL. After washing off the excess PLL in the same buffer, cells were swelled in hypotonic PHEM buffer (in mM: PIPES 60, HEPES 25, MgSO_4_ 4, EGTA 10, pH 6.96/KOH; 1 part of buffer mixed with 2 parts of deionized water) for 30 seconds. Unroofing was carried out in PHEM buffer using Branson 250 ultrasonifier with a 13 mm tip attached to the resonator at the lowest strength, pulse duration of 350-400 ms; the tip was placed at a distance of about 5 mm from the coverslip.

After unroofing, cells were fixed instantly or after a specified delay in PHEM buffer containing 1% glutaraldehyde (GA) (TEM grade, Serva GmbH, Germany) for 15 minutes at room temperature (RT). For the “delay experiments” for time point “0” the cells were unroofed in the PHEM buffer containing 0.2% GA and then fixed as it mentioned above; otherwise the cells were unroofed and incubated in PHEM buffer for 5, 10, 15 or 20 minutes at RT) followed by fixation with GA for 125 minutes at RT. Then, the samples were extensively washed with PHEM buffer and additionally fixed for 15 minutes in PHEM supplemented with tannic acid 0.2% (Sigma Aldrich, USA) followed by a thorough wash with deionized water. Then the samples were stabilized with aqueous FeCl_3_ 0.2% for 15 minutes. After washing off FeCl_3_ cells were dehydrated in ethanol series (50%, 70%, 2x 100%, 1 minute each, agitated), incubated with agitation for 1 minute in the 1:1 mixture of 100% ethanol with hexamethyldisilazane (HMDS; Carl Roth, Germany), transferred to pure HMDS (X2, 1 min each, agitated) and then air dried. Regions of coverslips containing unroofed Xenapses were visually identified using phase-contrast light microscopy, cut with a diamond cutter, glued on the aluminum sample carriers with CCC (‘Conductive Carbon Cement after Göcke’, Plano GmbH, Germany), and rotary coated after full drying of cement with Pt/C 2 nm at 45° in Leica ACE900. Scanning electron microscopy was performed with FEG SEM Hitachi S5000 (Hitachi, Japan) at 30 kV using a secondary electron detector; images were acquired and digitized using DISS5 (Point Electronic, Germany). The data analysis (CCS size and type distribution) was performed in MetaMorph (Molecular Devices, USA) and OriginLab (OriginLab Co, USA).

### Live TIRF microscopy

TIRF image acquisition was performed by a cooled sCMOS PRIME 95B camera (11 µm x 11 µm pixel area, Photometrics) controlled by μManager imaging software (μManager paper). The camera was side mounted onto the Nikon Eclipse Ti microscope equipped with a TIRF illumination module and 100x / 1.49 NA Apo TIRF oil immersion objective. Field stimulation was achieved using platinum electrodes and applying 80 mA pulse with alternating polarity administered by a constant current stimulus isolator (WPI A 385, World Precision Instruments). 50 µM D,L-2-amino-5-phosphonovaleric acid (AP5), and 10 µM 6-cyano-7-nitroquinoxaline-2,3-dione (CNQX) were added to the measuring buffer to prevent recurring neuronal activity. An OMICRON SOLE-6 laser light engine with high-speed analog and digital modulation controlled the lasers. The relevant fluorescent constructs used in this study were illuminated with either 488 nm, 638 nm (both diode lasers) or 561 nm (diode-pumped solid-state laser).

For photobleaching experiments, the same imaging conditions as for the stimulus assays were used and instead of the stimulus train a continuous 2 second FRAP pulse (100% laser intensity, 225 mW at 488 nm) was provided. mEos3.2-CLC was imaged by toggling the excitation lasers at 488 nm and 561 nm with a 100 ms interval and photoconversion was achieved with a 2 second 405 nm pulse (100% intensity, 180 mW). Measurement at 37°C was performed by placing the measuring chamber in the Heating insert and Incubator PM 2000 on the microscope mechanical stage, which in turn were connected to the Tempcontrol 37-2 Digital Microscope Temperature Process Controller (PeCon Gmbh).

An Arduino microcontroller unit (Arduino IDE 1.6.0) with a self-written code functioned as the external trigger controlling and synchronizing with high precision the lasers, camera, and stimulus isolator.

### STED and Confocal microscopy

Imaging with Leica TCS SP8 STED 3D microscope was controlled by LAS X software (Leica, Wetzlar, Germany). Cells expressing Halo-CLC (Fig. 1b) were live stained with membrane permeable JF646 (Promega, WI, USA) by incubating the cells with the dye (10 µM) in the preconditioned culture media for 10 min at 37°C. Cells were unroofed at 4°C and mounted immediately without further processing onto the glycerol immersion objective (93×/ 1.30 NA HC PL APO CORR) for imaging at room temperature. Excitation laser line 633 nm to image JF646 was generated by an 80 MHz tunable white light and a pulsed 775 nm depletion beam was used to create the xy donut. Images were acquired at 1 Hz and 0.1 Hz by sequential scanning performed in the resonance scanning mode with very similar parameters between the two acquisition rates to minimize differences in bleaching (36-38% STED laser intensity, line average of 8, 25 nm pixel size, 0.73 µs pixel dwell time). A gating of 0.5 – 6 ns was set before detection with HyD detectors.

The Leica setup was also employed for live Confocal imaging of EGFP-CLCa expressing Xenapses in Ringers buffer (Fig 2cd). The FRAP pulse lasting 0.7 s was provided using the FRAP booster and FRAP zoomer option to maximize bleaching within the ROI (100% intensity 500 mW). Imaging was performed at ∼ 3 % WLL intensity using the high resolution Galvo scanner mode with detection by a photomultiplier. 10 images, with pixel size of 150 nm, were acquired every 1s except for the image immediately after the FRAP pulse to account for the diffusion component.

STED imaging at room temperature was performed using a STEDYCON scanner (Abberior instruments GmbH, Göttingen, Germany) attached to a Nikon Ti-E microscope with cells mounted on a 100x / 1.45 NA oil immersion objective (Fig.3). Cells transfected with Halo-CLCa were live stained as described above. For live visualization of Clathrin coated structures (Fig. 3a), the medium with dye was washed off and replaced with Ringers buffer. For simultaneous visualization of endogenous Clathrin and overexpressed Halo-CLCa (Fig. 3d), Xenapses were unroofed and were fixed in 4% PFA in PHEM followed by staining with anti-Clathrin heavy chain primary antibody (Abcam, Cambridge, UK) and Alexa594 goat ant-rabbit (Invitrogen, MA, USA) secondary antibody for 15 minutes each at RT in blocking buffer (PHEM + 1.5 % BSA). The in-built STEDYCON Smart Control software was used for image acquisition. All imaging was done with a pixel dwell time of 10 µs and pixel size of 20 nm was achieved with 60-70% depletion laser intensity. Images were acquired with a 600 – 650 nm detector consisting of avalanche photodiodes with a time gating of 1 to 7 ns. Lasers at 640 nm and 561 nm were used to excite the JF646 dyes and AlexaFluor 594 respectively with the xy donut achieved with a 775 nm pulsed depletion laser.

### TIRF image analysis

All images were analyzed using self-written MATLAB (Mathworks, CA, USA) scripts. The recorded image stack was loaded onto MATLAB and their mean was generated. The individual Xenapse structures were identified by converting the greyscale image mean to binary mask using thresholding, a function which calculates the luminance threshold value in the range (0,1) and converts to a binary image. Whole Xenapse responses were extracted by applying the binary image mask on to the raw image. A MATLAB based graphical user interface (GUI) was used to select for Xenapses showing a stimulus-dependent response.

Clathrin stimulus-dependent response curves were identified by combining the thresholding derived mask with an additional binary mask derived from an à trous wavelet transformation of the baseline average ^9^. The mask was applied onto the raw image stack and the responding Xenapses were selected using the GUI as mentioned above.

The selected structures were further processed to derive sub-Xenapse structures of maximal endocytic activity used to derive the final response curve. This was achieved by a binary mask-based difference images of the baseline (20 frames before stimulus) and back baseline (final 10 frames) with the frames around the endocytic minimum respectively.

The FRAP responses curves were derived as for the Clathrin and AP2 stimulus-dependent except no GUI-based selection was applied. The final FRAP traces are based on sub-Xenapse structures generated from the difference image of the10 final frames with the frame 3 seconds after the bleach minimum. The resulting substructures were identified as areas of maximum post-photobleaching recovery with minimized diffusional component.

### STED and Confocal image analysis

All images were analyzed using self-written MATLAB (Mathworks) script. For the analysis of Clathrin dynamics in unroofed Xenapses (Fig. 1b) a PSF based localization was performed on the averaged image of the 10 STED image stack to identify single CCSs. A 7 × 7 pixel cut out (0.031 µm^2^) was made for each localization and applied onto the image stack to generate the fluorescence trace.

For the confocal FRAP data (Fig. 2cd), the FRAP region was derived from the difference of the pre-FRAP baseline average (images 1-3) and the FRAP frame (image 4, acquired immediately after the photobleaching pulse). A binary mask derived from the à trous wavelet transformation of the difference image was applied on the raw image stack to generate the FRAP trace. The non-FRAP region was identified from the threshold adjusted binary mask of the FRAP frame (image 4) which contains the pixels not subjected to the photobleaching pulse.

Aberrior STED images acquired at 20 s interval (Fig. 3a) were processed by ImageJ to correct for any drift and normalize for difference in intensities between images resulting from photobleaching (Process>Enhance Contrast>Normalize). Rolling ball radius of 10 pixels was used further to reduce background. The processed images were transferred to MATLAB and converted to binary images based on thresholding. Connected components smaller than 5 pixels (pixel size 20 nm) in size were removed from the binary images as these were deemed to be derived from background noise resulting from non-specific dye staining (*bwareaopen*). Multiplying one binary image by a factor of 2 and adding the second image allowed discrimination of Clathrin sub-structures that were unique to one of the two time points or co-occured at both time points (upper row). Connected components were determined for the respective binary images and filtered by properties area, ellipticity and solidity (*bwconncomp, regionprops*) to detect bonafide Clathrin pits (lower row).

### Quantification of CCSs in SEM

SEM images were analyzed manually in MetaMorph based on a previously published approach ^30^. First, the area of each unroofed Xenapse was segmented by outlining the edge of the remaining membrane. If the footprint of the Xenapse was bigger than the opening, only the exposed surface was taken for the area estimation. Next, the following categories were used for the quantitative analysis of the CCSs, namely: ‘initial structure’ – the CCS that consist of one to three penta-or hexagons and has the open ends, ‘lattice’ – the flat Clathrin structure with no visible curvature, ‘domed’ – visibly curved structure with open ends and ‘pit’ – fully formed spheres with no open ends visible. The latter structures were also much brighter at the circumference due to higher yield of secondary electrons. Then all the CCSs per Xenapse were counted and the fraction of each category was determined. The diameter of the domed and pit CCSs were determined by simplifying that the structures were of spherical shape; the area was calculated only as the area of the visible portion of the dome or the pit, as the real curvature could not be determined from SEM images (thus the area of the CCSs is greatly underestimated).

## Acknowledgements

We thank K. Tkotz and A. Roetrige for excellent technical support, and J. Duan for help with the Xenapse preparation. We also thank M.Kahms for proofreading the manuscript. This work was supported by grants of the Deutsche Forschungsgemeinschaft (SFB 944 and SFB 1348) to J.K.

## Author information

The authors declare no competing financial interests. Correspondence and requests for materials should be addressed to J.K.

## Author contributions

S.K., Y.T. and J.K. conceptualized the study and designed the experiments; S.K. conducted the imaging experiments; S.K. and J.K. analyzed the imaging data; Y.T and N.G conducted and analyzed EM experiments, J.L. wrote software for data analysis; S.K. and J.K. wrote the manuscript.

## Data availability

The data that support the findings of this study are available from the corresponding author upon reasonable request.

## Code availability

Matlab Code used for analyzing the data presented in this manuscript is available from the corresponding author upon request.

---

**Extended Data Fig. 1.**
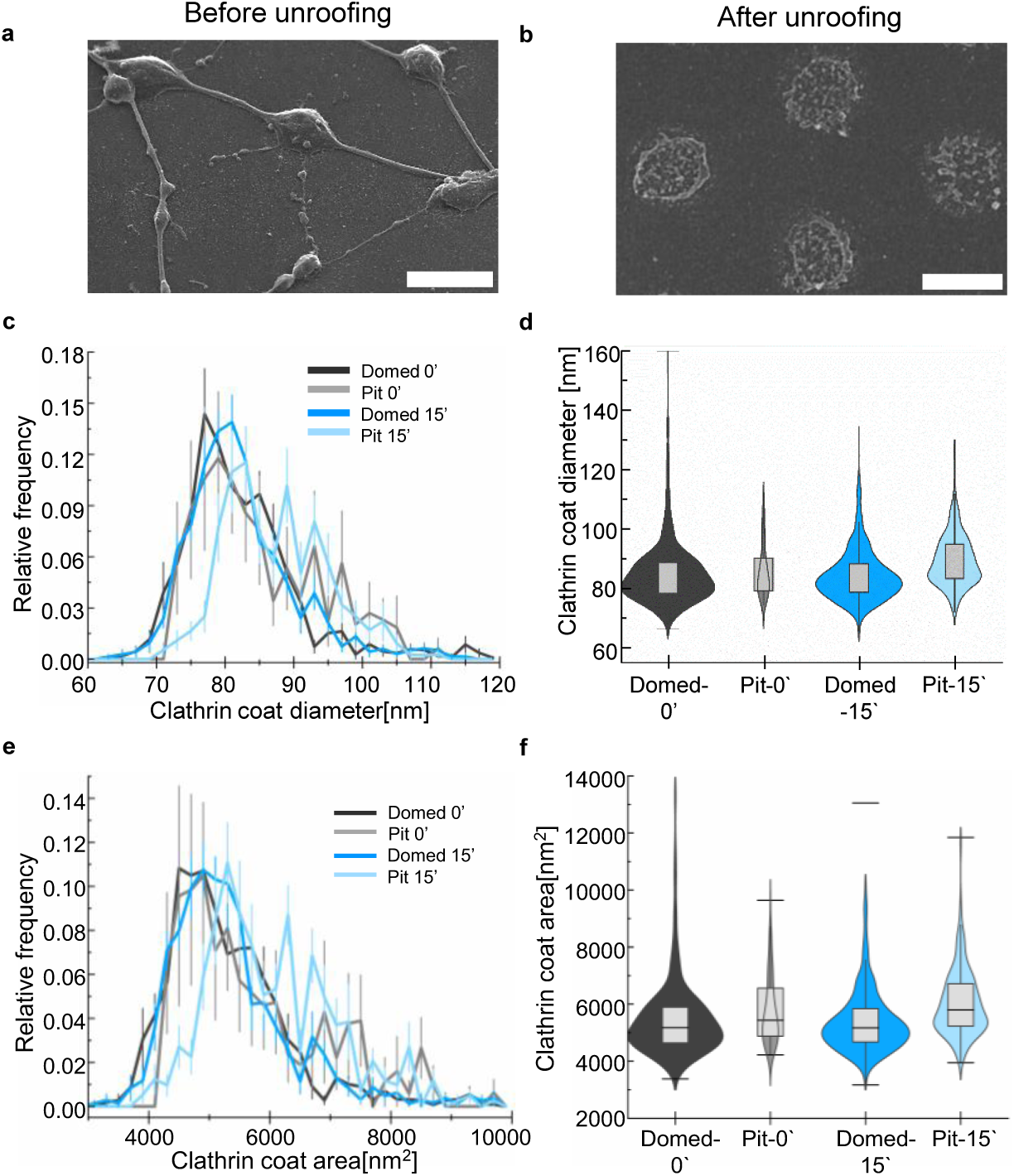
Generation and unroofing of Xenapses. SEM images of Xenapses before unroofing with the intact apical membranes (**a**) and after unroofing (**b**). Scale bar represents 4 µm. **c**, Size distribution of domed and pit-like CCS in unroofed xenapses fixed at 0 and 15 min. after unroofing represented as the diameter of CCS estimated from 2D images. **d**, Box-chart of the diameter of CCS representing all data in (**b**) with the outliers included. **e**, Size distribution of domed and pit-like CCS in unroofed xenapses fixed at 0 and 15 min after unroofing represented as the area of CCS estimated from 2D images. **f**, Box-chart of the area of CCS representing all data in (**d)** with the outliers included.

**Extended Data Fig. 2.**
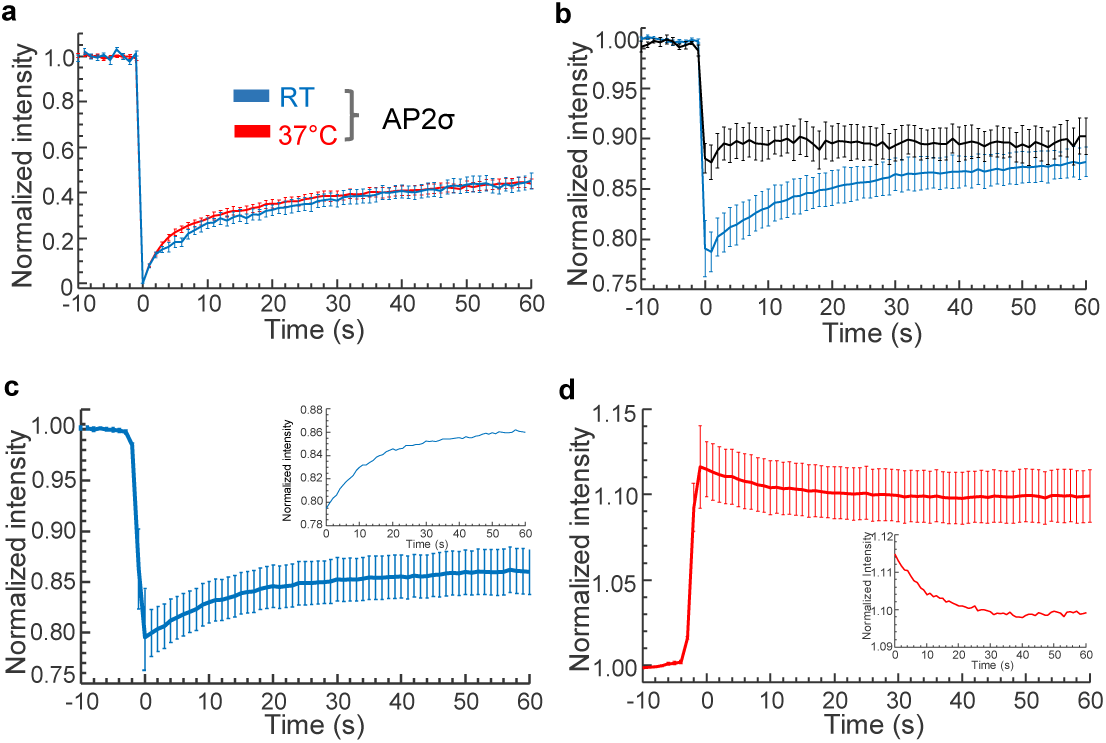
EW-FRAP response curves of fluorescently labelled endocytic proteins. **a**, Normalized average of AP2σ-EGFP fluorescence recovery traces at RT (n = 9 measurements) and 37°C (n = 14 measurements) in response to a TIRF photobleaching pulse revealing the amplitude of recovery. **b**, Normalized average fluorescence recovery traces of à trous wavelet-filtered boutons at RT of Dynamin 1-EGFP (n = 8 measurements) and EGFP-CLCa from Fig.3C. **c,d**, Normalized average fluorescence trace at RT of Xenapses expressing mEos3.2-CLCa with a photoconversion pulse at 405 nm revealing the recovery (**c**, 488 nm) and removal (**d**, 561 nm) of surface Clathrin in the same boutons (n = 10 measurements). The insets show the respective traces without the baseline and a scaled down y-axis. All traces are shown as mean ± s.e.m.

**Extended Data Fig. 3.**
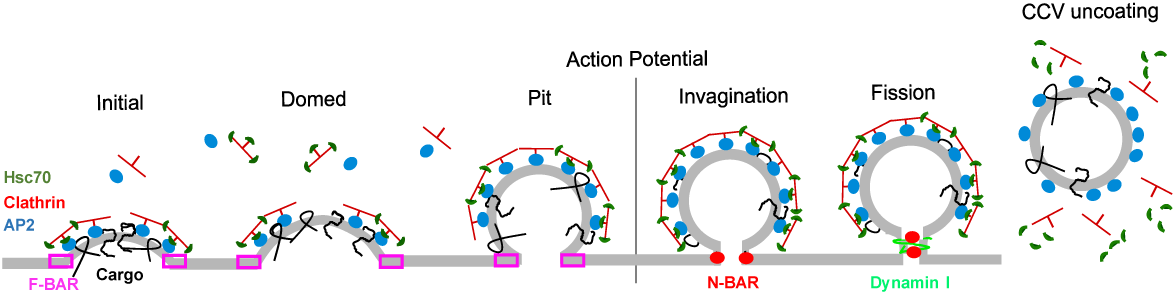
Schematic summary of RRetP life cycle before and after action potential. Schematic of the organisation of readily retrievable pool (RRetP) and the accessory proteins. At steady state the Clathrin coated RRetP dynamically changes in the degree of assembly and curvature. F-BAR proteins contribute to the initial curvature but fluctuation between different RRetP states is driven by Hsc70-mediated Clathrin exchange and the associated exchange of the adaptor AP2. Following SV fusion in response to an action potential, the pit structures predominate with the N-BAR protein mediated invagination and the ensuing recruitment of Dynamin 1 resulting in scission. The Clathrin-coated vesicle subsequently undergoes Hsc70 mediated uncoating.

## References

1. Roth, T. F. & Porter, K. R. Yolk protein uptake in the oocyte of the mosquito Aedes Aegypti. L. J Cell Biol 20, 313–332; 10.1083/jcb.20.2.313 (1964).

2. Kanaseki, T. & Kadota, K. The “vesicle in a basket”. A morphological study of the coated vesicle isolated from the nerve endings of the guinea pig brain, with special reference to the mechanism of membrane movements. J Cell Biol 42, 202–220; 10.1083/jcb.42.1.202 (1969).

3. Pearse, B. M. Coated vesicles from pig brain: purification and biochemical characterization. Journal of molecular biology 97, 93–98; 10.1016/s0022-2836(75)80024-6 (1975).

4. Ungewickell, E. & Branton, D. Assembly units of clathrin coats. Nature 289, 420–422; 10.1038/289420a0 (1981).

5. Motley, A., Bright, N. A., Seaman, M. N. J. & Robinson, M. S. Clathrin-mediated endocytosis in AP-2-depleted cells. J Cell Biol 162, 909–918; 10.1083/jcb.200305145 (2003).

6. Chen, Z. & Schmid, S. L. Evolving models for assembling and shaping clathrin-coated pits. J. Cell Biol. 219; 10.1083/jcb.202005126 (2020).

7. Pascolutti, R. et al. Molecularly Distinct Clathrin-Coated Pits Differentially Impact EGFR Fate and Signaling. CELL REPORTS 27, 3049–3061.e6; 10.1016/j.celrep.2019.05.017 (2019).

8. Heuser, J. Three-dimensional visualization of coated vesicle formation in fibroblasts. J Cell Biol 84, 560–583; 10.1083/jcb.84.3.560 (1980).

9. Lehrich J, Glyvuk N, Duan J, Krishnan S, Tsytsyura Y, Keller U, Penothil Sunil L, Nosov G, Hüve J, Selenschik P, Reißner C, Missler M, Kahms M, Klingauf J. Optical recordings of synaptic vesicle fusion reveal diffusional dispersion. Biorxiv. (2025).

10. Krishnan, S. & Klingauf, J. The readily retrievable pool of synaptic vesicles. Biological chemistry 404, 385–397; 10.1515/hsz-2022-0298 (2023).

11. Wienisch, M. & Klingauf, J. Vesicular proteins exocytosed and subsequently retrieved by compensatory endocytosis are nonidentical. Nat. Neurosci. 9, 1019–1027; 10.1038/nn1739 (2006).

12. Mueller, V. J., Wienisch, M., Nehring, R. B. & Klingauf, J. Monitoring clathrin-mediated endocytosis during synaptic activity. The Journal of neuroscience: the official journal of the Society for Neuroscience 24, 2004–2012; 10.1523/JNEUROSCI.4080-03.2004 (2004).

13. Hua, Y. et al. A readily retrievable pool of synaptic vesicles. Nature Neuroscience 14, 833–U42; 10.1038/nn.2838 (2011).

14. Avinoam, O., Schorb, M., Beese, C. J., Briggs, J. A. G. & Kaksonen, M. ENDOCYTOSIS. Endocytic sites mature by continuous bending and remodeling of the clathrin coat. Science (New York, N.Y.) 348, 1369–1372; 10.1126/science.aaa9555 (2015).

15. Loerke, D., Wienisch, M., Kochubey, O. & Klingauf, J. Differential control of clathrin subunit dynamics measured with EW-FRAP microscopy. Traffic 6, 918–929; 10.1111/j.1600-0854.2005.00329.x (2005).

16. Wu, X. et al. Clathrin exchange during clathrin-mediated endocytosis. J Cell Biol 155, 291–300; 10.1083/jcb.200104085 (2001).

17. Nossal, R. & Zimmerberg, J. Endocytosis: curvature to the ENTH degree. Current Biology 12, R770–2; 10.1016/s0960-9822(02)01289-7 (2002).

18. Kirchhausen, T. Coated pits and coated vesicles — sorting it all out. Current Opinion in Structural Biology 3, 182–188; 10.1016/S0959-440X(05)80150-2 (1993).

19. Saffarian, S., Cocucci, E. & Kirchhausen, T. Distinct dynamics of endocytic clathrin-coated pits and coated plaques. Plos Biology 7, e1000191; 10.1371/journal.pbio.1000191 (2009).

20. Willy, N. M. et al. De novo endocytic clathrin coats develop curvature at early stages of their formation. Developmental cell 56, 3146–3159.e5; 10.1016/j.devcel.2021.10.019 (2021).

21. Bucher, D. et al. Clathrin-adaptor ratio and membrane tension regulate the flat-to-curved transition of the clathrin coat during endocytosis. Nature Communications 9, 1109; 10.1038/s41467-018-03533-0 (2018).

22. Heuser, J. E. & Anderson, R. G. Hypertonic media inhibit receptor-mediated endocytosis by blocking clathrin-coated pit formation. J Cell Biol 108, 389–400; 10.1083/jcb.108.2.389 (1989).

23. Mund, M. et al. Clathrin coats partially preassemble and subsequently bend during endocytosis. J. Cell Biol. 222; 10.1083/jcb.202206038 (2023).

24. Watanabe, S. et al. Ultrafast endocytosis at mouse hippocampal synapses. Nature 504, 242–247; 10.1038/nature12809 (2013).

25. Imoto, Y. et al. Dynamin is primed at endocytic sites for ultrafast endocytosis. Neuron 110, 2815–2835.e13; 10.1016/j.neuron.2022.06.010 (2022).

26. Delvendahl, I., Vyleta, N. P., Gersdorff, H. von & Hallermann, S. Fast, Temperature-Sensitive and Clathrin-Independent Endocytosis at Central Synapses. Neuron 90, 492– 498; 10.1016/j.neuron.2016.03.013 (2016).

27. Vacquier, V. D. The isolation of intact cortical granules from sea urchin eggs: calcium lons trigger granule discharge. Developmental biology 43, 62–74; 10.1016/0012-1606(75)90131-1 (1975).

28. Clarke, M., Schatten, G., Mazia, D. & Spudich, J. A. Visualization of actin fibers associated with the cell membrane in amoebae of Dictyostelium discoideum. PNAS 72, 1758–1762; 10.1073/pnas.72.5.1758 (1975).

29. Heuser, J. Effects of cytoplasmic acidification on clathrin lattice morphology. Journal of Cell Biology 108, 401–411 (1989).

30. Sochacki, K. A. et al. The structure and spontaneous curvature of clathrin lattices at the plasma membrane. Developmental cell 56, 1131–1146.e3; 10.1016/j.devcel.2021.03.017 (2021).

31. Vassilopoulos, S. & Montagnac, G. Clathrin assemblies at a glance. Journal of cell science 137; 10.1242/jcs.261674 (2024).

32. Grove, J. et al. Flat clathrin lattices: stable features of the plasma membrane. Molecular Biology of the Cell 25, 3581–3594; 10.1091/mbc.E14-06-1154 (2014).

33. Xing, Y. et al. Structure of clathrin coat with bound Hsc70 and auxilin: mechanism of Hsc70-facilitated disassembly. EMBO J. 29, 655–665; 10.1038/emboj.2009.383 (2010).

34. Beck, K. A. & Keen, J. H. Interaction of phosphoinositide cycle intermediates with the plasma membrane-associated clathrin assembly protein AP-2. The Journal of biological chemistry 266, 4442–4447 (1991).

35. Krishnan, S., Lehrich, J., Tsytsyura, Y., Glyvuk, N., Duan, J., Klingauf, J. Slow scission of single synaptic vesicles by Dynamin at physiological temperature. Biorxiv. 10.1101/2025.03.18.643914 (2025).

36. Jiang, R., Gao, B., Prasad, K., Greene, L. E. & Eisenberg, E. Hsc70 chaperones clathrin and primes it to interact with vesicle membranes. The Journal of biological chemistry 275, 8439–8447; 10.1074/jbc.275.12.8439 (2000).

37. Yim, Y.-I. et al. Exchange of clathrin, AP2 and epsin on clathrin-coated pits in permeabilized tissue culture cells. Journal of cell science 118, 2405–2413; 10.1242/jcs.02356 (2005).

38. Eisenberg, E. & Greene, L. E. Multiple roles of auxilin and hsc70 in clathrin-mediated endocytosis. Traffic 8, 640–646; 10.1111/j.1600-0854.2007.00568.x (2007).

39. Chen, Y. et al. Dynamic instability of clathrin assembly provides proofreading control for endocytosis. J. Cell Biol. 218, 3200–3211; 10.1083/jcb.201804136 (2019).

40. Ungewickell, E. The 70-kd mammalian heat shock proteins are structurally and functionally related to the uncoating protein that releases clathrin triskelia from coated vesicles. EMBO J. 4, 3385–3391; 10.1002/j.1460-2075.1985.tb04094.x (1985).

41. Morgan, J. R., Prasad, K., Jin, S., Augustine, G. J. & Lafer, E. M. Uncoating of clathrin-coated vesicles in presynaptic terminals: roles for Hsc70 and auxilin. Neuron 32, 289– 300; 10.1016/s0896-6273(01)00467-6 (2001).

42. Rapoport, I., Boll, W., Yu, A., Böcking, T. & Kirchhausen, T. A motif in the clathrin heavy chain required for the Hsc70/auxilin uncoating reaction. Molecular Biology of the Cell 19, 405–413; 10.1091/mbc.e07-09-0870 (2008).

43. Fourie, A. M., Sambrook, J. F. & Gething, M. J. Common and divergent peptide binding specificities of hsp70 molecular chaperones. The Journal of biological chemistry 269, 30470–30478 (1994).

44. Takenaka, I. M., Leung, S. M., McAndrew, S. J., Brown, J. P. & Hightower, L. E. Hsc70-binding peptides selected from a phage display peptide library that resemble organellar targeting sequences. The Journal of biological chemistry 270, 19839–19844; 10.1074/jbc.270.34.19839 (1995).

45. Newmyer, S. L., Christensen, A. & Sever, S. Auxilin-dynamin interactions link the uncoating ATPase chaperone machinery with vesicle formation. Developmental cell 4, 929–940; 10.1016/s1534-5807(03)00157-6 (2003).

46. Yim, Y.-I. et al. Endocytosis and clathrin-uncoating defects at synapses of auxilin knockout mice. Proceedings of the National Academy of Sciences 107, 4412–4417; 10.1073/pnas.1000738107 (2010).

47. Hirst, J. et al. Auxilin depletion causes self-assembly of clathrin into membraneless cages in vivo. Traffic 9, 1354–1371; 10.1111/j.1600-0854.2008.00764.x (2008).

48. Jin, A. J. & Nossal, R. Rigidity of triskelion arms and clathrin nets. Biophysical journal 78, 1183–1194; 10.1016/S0006-3495(00)76676-8 (2000).

49. Mashl, R. J. & Bruinsma, R. F. Spontaneous-curvature theory of clathrin-coated membranes. Biophysical journal 74, 2862–2875; 10.1016/S0006-3495(98)77993-7 (1998).

50. Dannhauser, P. N. & Ungewickell, E. J. Reconstitution of clathrin-coated bud and vesicle formation with minimal components. Nature cell biology 14, 634–639; 10.1038/ncb2478 (2012).

51. Nawara, T. J. et al. Imaging vesicle formation dynamics supports the flexible model of clathrin-mediated endocytosis. Nature Communications 13, 1732; 10.1038/s41467-022-29317-1 (2022).

52. Zeno, W. F. et al. Clathrin senses membrane curvature. Biophysical journal 120, 818– 828; 10.1016/j.bpj.2020.12.035 (2021).

53. Morris, K. L. et al. Cryo-EM of multiple cage architectures reveals a universal mode of clathrin self-assembly. Nature structural & molecular biology 26, 890–898; 10.1038/s41594-019-0292-0 (2019).

54. He, K. et al. Dynamics of Auxilin 1 and GAK in clathrin-mediated traffic. J. Cell Biol. 219; 10.1083/jcb.201908142 (2020).

55. Foss, S. M., Li, H., Santos, M. S., Edwards, R. H. & Voglmaier, S. M. Multiple dileucine-like motifs direct VGLUT1 trafficking. The Journal of Neuroscience 33, 10647–10660; 10.1523/JNEUROSCI.5662-12.2013 (2013).

56. Kovtun, O., Dickson, V. K., Kelly, B. T., Owen, D. J. & Briggs, J. A. G. Architecture of the AP2/clathrin coat on the membranes of clathrin-coated vesicles. Science advances 6, eaba8381; 10.1126/sciadv.aba8381 (2020).

57. Voglmaier, S. M. et al. Distinct endocytic pathways control the rate and extent of synaptic vesicle protein recycling. Neuron 51, 71–84; 10.1016/j.neuron.2006.05.027 (2006).

58. Heuser, J. The production of ‘cell cortices’ for light and electron microscopy. Traffic 1, 545–552; 10.1034/j.1600-0854.2000.010704.x (2000).

59. Sochacki, K. A., Shtengel, G., van Engelenburg, S. B., Hess, H. F. & Taraska, J. W. Correlative super-resolution fluorescence and metal-replica transmission electron microscopy. Nat Methods 11, 305–308; 10.1038/nmeth.2816 (2014).

